# Facemap: a framework for modeling neural activity based on orofacial tracking

**DOI:** 10.1101/2022.11.03.515121

**Authors:** Atika Syeda, Lin Zhong, Renee Tung, Will Long, Marius Pachitariu, Carsen Stringer

**Affiliations:** HHMI Janelia Research Campus; Johns Hopkins University

## Abstract

Recent studies in mice have shown that orofacial behaviors drive a large fraction of neural activity across the brain. To understand the nature and function of these signals, we need better computational models to characterize the behaviors and relate them to neural activity. Here we developed Facemap, a framework consisting of a keypoint tracking algorithm and a deep neural network encoder for predicting neural activity. We used the Facemap keypoints as input for the deep neural network to predict the activity of ∼50,000 simultaneously-recorded neurons and in visual cortex we doubled the amount of explained variance compared to previous methods. Our keypoint tracking algorithm was more accurate than existing pose estimation tools, while the inference speed was several times faster, making it a powerful tool for closed-loop behavioral experiments. The Facemap tracker was easy to adapt to data from new labs, requiring as few as 10 annotated frames for near-optimal performance. We used Facemap to find that the neuronal activity clusters which were highly driven by behaviors were more spatially spread-out across cortex. We also found that the deep keypoint features inferred by the model had time-asymmetrical state dynamics that were not apparent in the raw keypoint data. In summary, Facemap provides a stepping stone towards understanding the function of the brainwide neural signals and their relation to behavior.

## Introduction

Neurons across the brain are constantly active, even in the absence of external sensory stimuli or a behavioral task [1, 2]. This ongoing, spontaneous neural activity is driven by the spontaneous behaviors of the animal, such as running, head movements and whisking in mice [3–9], tail movements in zebrafish [10], and body movements in flies [11–13]. In mice, different neurons were best explained by different combinations of orofacial behaviors, such as whisking, sniffing and grooming, showing that multi-dimensional representations of behavior exist across the brain [14– 17]. These multi-dimensional behavioral representations persist during presentations of sensory stimuli [14] and during decision-making tasks [18–20]. Despite the widespread presence of behavioral signals across the brain, their role and function remains poorly understood. To make progress on understanding these neural signals, it is important to develop better computational models. This requires progress in two areas: 1) better quantification of orofacial behavior, 2) better models of the influence of behavior on neural activity.

To quantify behavior, previous studies took advantage of the stability of the head-fixed experimental setup to compute low-dimensional features of the raw behavior movies, either using principal components of the movies [14, 17, 20], or using autoencoders fit to the movies [21, 22]. Although movie principal components are easy to compute, the resulting features are hard to interpret. Another common approach for quantifying orofacial movements is whisker tracking, which can provide specific and interpretable information about whisker motion [23–26]. However, previous approaches for whisker tracking required trimming the other whiskers and/or whisker painting, which may alter mouse behavior, and they also required a high-speed overhead camera, which may be unavailable in many experimental setups. An alternative approach is markerless pose estimation, or keypoint tracking. Several algorithms exist for general keypoint tracking in animals [27–31], but none of these tools have specialized methods for tracking orofacial movements.

Similarly, better models are needed to account for the influence of behavior on neural activity. Previous studies used simple approaches like reduced-rank regression or ridge regression [14, 20]. These models are linear and do not take into account temporal dynamics. They are therefore unlikely to capture the full influence of time-varying, multi-dimensional behavior on neural activity.

To address these shortcomings, we developed Facemap, a framework that quantifies orofacial behaviors using keypoint tracking and uses those key-points to predict neural activity. Both steps of the framework are powered by deep neural networks. To track orofacial behaviors, we developed a pose estimation tool that tracks 13 distinct keypoints on the mouse face from variable camera views. Our pose estimation tool is more accurate than the best existing method (DeepLabCut), and it is also twice as fast, thus providing a viable option for online behavioral tracking. On new data, the Facemap tracker requires only 10 new labelled frames for near-optimal performance. We also developed a multi-layer neural network that is optimized to predict neural dynamics from keypoint dynamics. Compared to previous methods, this approach can predict almost twice as much neural variance for neurons in visual cortex. Furthermore, the model learns deep keypoint features that have highly structured state dynamics, which we inferred using a Hidden Markov Model. Hence, Facemap can be used to obtain insights into both the structure and influence of orofacial behaviors on neural activity, thus providing a stepping stone towards understanding the function of the brainwide behavioral signals.

## Results

### Fast and accurate tracker for mouse orofacial movements

We start by describing a neural network model for key-point tracking on the mouse face, the Facemap tracker. As a first step, we chose several well-defined keypoints that could track various orofacial movements (Figure 1a). To capture whisking, we tracked three whiskers that are visible from most camera views, labeling the points at the base of the whiskers. To capture sniffing, we tracked four nose-related keypoints (bottom, top, tip, and right-bottom, when in view). To capture mouth movements, when the mouth was in view, we tracked two mouth keypoints (mouth and lower lip). To capture eye movements, such as blinking, we tracked the four corners of the eye (bottom, top, front, back). We did not track the pupil, because it is completely dilated and un-trackable in darkness, and also because it is easier to track with simpler methods [14].

**Figure 1:**
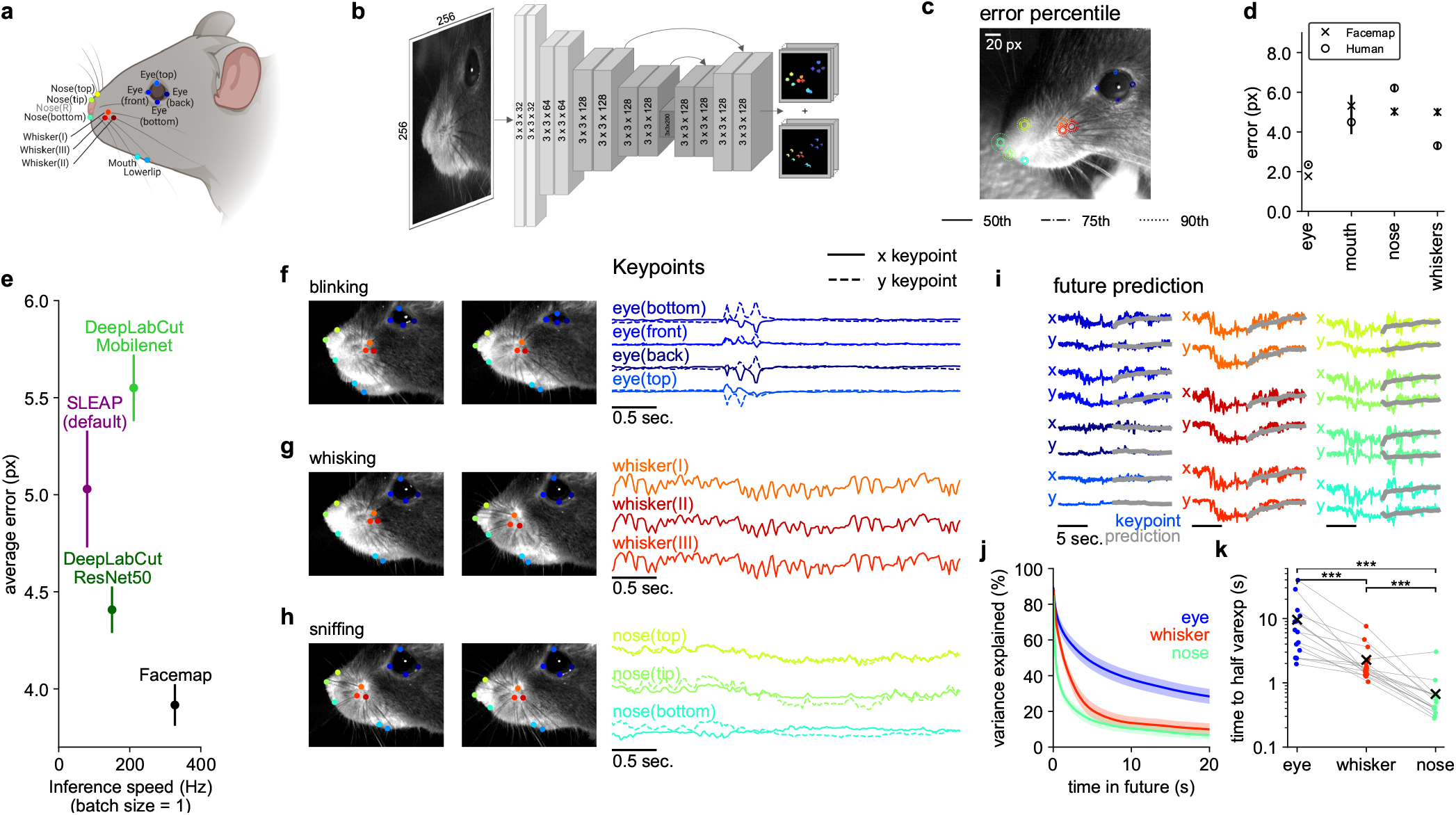
Fast and accurate mouse orofacial keypoint tracking. **a**, 13 distinct keypoints selected for tracking the eye, mouth, whiskers and nose on the mouse face. **b**, Architecture of the Facemap network, a U-Net style convolutional neural network. **c**, The error percentiles across test frames from a new mouse, where error is defined as the Euclidean distance between the ground-truth label and the prediction. **d**, Summary of Facemap performance on test data for different subgroups of keypoints. Human error shown for a subset of the test frames labeled in two different sessions by a human annotator. Error bars represent s.e.m., n=400, 95, 361 and 300 keypoint labels for eye, mouth, nose and whiskers respectively across 100 test frames. **e**, The average error and inference speed of the Facemap tracker compared with other pose estimation tools. Error bars represent s.e.m., n=1156 keypoint labels. **f**, Traces of *x*- and *y*-coordinates of keypoints during different orofacial behaviors. **g**, Prediction of keypoint traces into the future (test data). **h**, Variance explained of future prediction at different time lags, summarized for each face region. Error bars represent s.e.m., n=16 recordings. **i**, Decay time to 50% of variance explained at 20ms timelag. The “x” represents the average. Two-sided Wilcoxon signed-rank test, *** denotes *p <* 0.001.

Our goal was to build a model that would generalize well to new data. To achieve this goal, we collected a dataset of short mouse face videos from many different mice with the camera set up at several different angles. From this dataset of 16 mice and 53 video recordings at different views, we manually annotated 2500 frames (Figure S1). We used 2400 frames for training the network and set aside a test set consisting of 100 frames from multiple views of a new mouse.

Unlike more general approaches, like DeepLabCut and SLEAP [27, 31], we only require our tracker to perform well on specific keypoints from the mouse face. Thus, we hypothesized that a minimal “U-Net”-style neural network [32] would be sufficient for the task while providing faster tracking compared to the existing, bigger models (Table S1). Similar to DeepLab-Cut, which in turn is based on DeeperCut [33], the Facemap tracker takes as input an image and outputs a set of downsampled probability heatmaps and location refinement maps to predict the *x* and *y* coordinates for each keypoint (Figure 1b). The likelihood values of the model prediction were used to filter the traces and remove outliers ([27], see Methods). The Facemap tracker was implemented from scratch using the neural network software PyTorch [34], which is a popular and easy-to-use alternative to the TensorFlow framework [35] used by DeepLabCut and SLEAP.

The keypoint error percentiles shown on an example test frame demonstrate the accuracy of the tracking (Figure 1c, Supplementary Video 1). To get an upper bound on the tracking performance, we manually labeled test frames twice at different orientations and compared the two sets of labels. We found that the tracker achieved human-level performance (Figure 1d). We compared our model with current state-of-the-art tools for keypoint tracking, DeepLab-Cut and SLEAP [27, 31, 36]. The Facemap tracker was more accurate than the other well-performing network, DeepLabCut with the ResNet50 backbone, both in average error (3.9 versus 4.4 pixels) and for individual keypoints (Figure 1e, Figure S2a). Facemap also out-performed DeepLabCut with the Mobilenet backbone, SLEAP default and SLEAP’s larger network (32 channels), which had average errors of 5.6, 5.0, and 5.7 pixels respectively (Figure 1e, Figure S2a).

To compare the speed of the networks for the purpose of online tracking, we computed the inference speed using a batch size of 1 (Figure 1e). All the networks can run faster with larger batch sizes, but only can a batch size of 1 be used for online inference of keypoints for closed-loop experiments. The smaller size of the Facemap tracker network provided a much faster inference speed of 327 Hz on a V100 GPU compared to DeepLabCut’s ResNet50 network (150 Hz), DeepLabCut’s Mobilenet network (211 Hz), SLEAP’s default network (80 Hz) and SLEAP’s larger network (c=32) (72 Hz). Across different GPU types Facemap consistently demonstrated the fastest inference speed (Table S2). We also benchmarked the inference speed of the Facemap tracker at larger batch sizes, and found that it was as fast or faster than all other networks across GPUs except for the Tesla T4 GPU, where DeepLabCut Mobilenet was fastest (Figure S2b). Therefore, Facemap is the fastest tracker with state-of-the-art performance, which enables its use in closed-loop experiments with high frame rates. The keypoints tracked by Facemap captured recognizable orofacial behaviors, such as blinking (Figure 1f), whisking (Figure 1g) and sniffing (Figure 1h), in addition to other orofacial behaviors. In the neural recordings, the camera view in Figure 1c was used, so mouth keypoints were not included in the analyses as they were not visible. Therefore, for the rest of this study, we use the eye, whisker and nose keypoints to characterize aspects of behavior and neural activity. To start, we investigated the timescales of the orofacial keypoints. To do this, we built a generalized linear model to predict the position of each keypoint in the future (prediction shown in Figure 1i). The variance explained of the model on test data decayed as a function of time into the future (Figure 1j). The predictability of the nose keypoints decayed fastest (∼1 second), followed by the whiskers (∼ 3 seconds) and eye keypoints (∼10 seconds) (Figure 1k). This was surprising because whisking was the fastest behavior observed in the videos (∼10Hz). However, these fast movements were pseudo-random (Figure 1g) and hard to predict and so they did not contribute strongly to the predictability of the whisker keypoints.

### Fine-tuning the Facemap tracker on new videos

We built the Facemap tracker to perform well on a variety of camera angles and across different mice. While Facemap generalized well on data from similar mice and camera configurations, the tracker had variable performance on videos from other labs (Figure 2a). We investigated whether a fine-tuning strategy might improve the performance of the tracker further on new data. We annotated a small set of video frames contributed by other labs to fine-tune the neural network individually for each lab. The fine-tuned network showed a dramatic drop in error after training with just one frame. Training with around 10 frames led to near-optimal performance (Figure 2b, Supplementary Videos 2).

**Figure 2:**
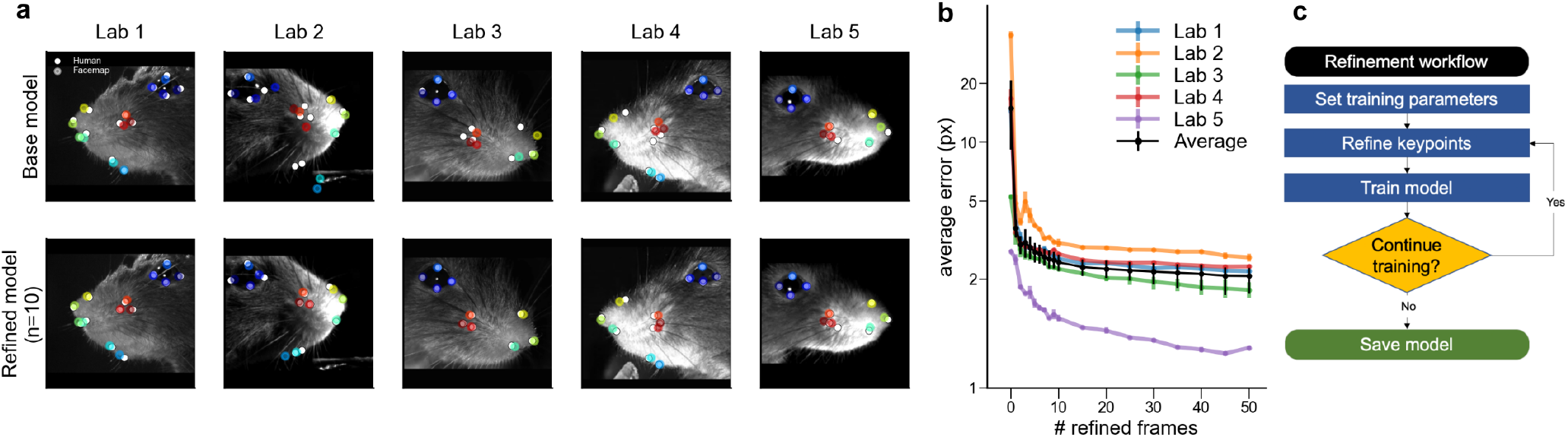
Keypoint tracking on mice from other labs by fine-tuning of the Facemap tracker. **a**, Top: Keypoint predictions using the Facemap tracker’s base model (white circles) and human annotations (colored circles) on mice from new experimental setups. Bottom: Keypoint predictions from the fine-tuned model trained with # refined frames = 10. **b**, Performance of the Facemap tracker measured by average error (px) on test frames as a function of the number of refined frames used for fine-tuning the base model (# refined frames=0 is the base model), for each lab and average test error across labs [black]. **c**, A flowchart of the refinement workflow implemented in our GUI.

Given the quick improvement in the network performance after fine-tuning, we reasoned this step is necessary for adapting Facemap’s tracker to new data. Therefore, we implemented a “human-in-the-loop” workflow to allow users to easily fine-tune Facemap for their own datasets in our graphical user interface (GUI) (Figure 2c). In the first step, the existing Facemap network is used to generate keypoint predictions. Next, the user refines the predicted keypoints to generate new training labels for the network. Then, the network is re-trained with the new labels to create a fine-tuned network. The fine-tuned network is then applied to new frames and the user can decide whether or not to repeat the retraining process depending on the performance of the fine-tuned network. Once a well-performing network is obtained, the fine-tuned network is saved in the GUI for future use. This fine-tuning step was also used for our experiments in the next section, where we combine keypoint tracking and large-scale neural recordings.

### A deep keypoint model of neural activity

To determine how neural activity depends on keypoint dynamics, we designed a neural network encoding model that can extract deep features from the keypoint data. Similar to deep encoding models in sensory neuroscience, the model has a first linear step for applying spatiotemporal filters to the time-varying keypoints (Figure 3a,b), followed by several fully-connected layers that process these features further, into more abstract representations that can better predict the neural activity. This deep keypoint model was trained end-to-end to predict the activity of the top 128 principal components of neural data from either visual or sensorimotor cortices (Figure 3c). The neural activity was split into training and test data in blocks of around 10 minutes and 3.5 minutes respectively and the variance explained was computed on the test data periods. We normalized the variance explained by an approximate upper-bound, estimated using peer prediction, similar to [14]. Compared to variations of the neural network architecture, the model we chose (Figure 3a) had the best performance while using the fewest number of layers (Figure S3).

**Figure 3:**
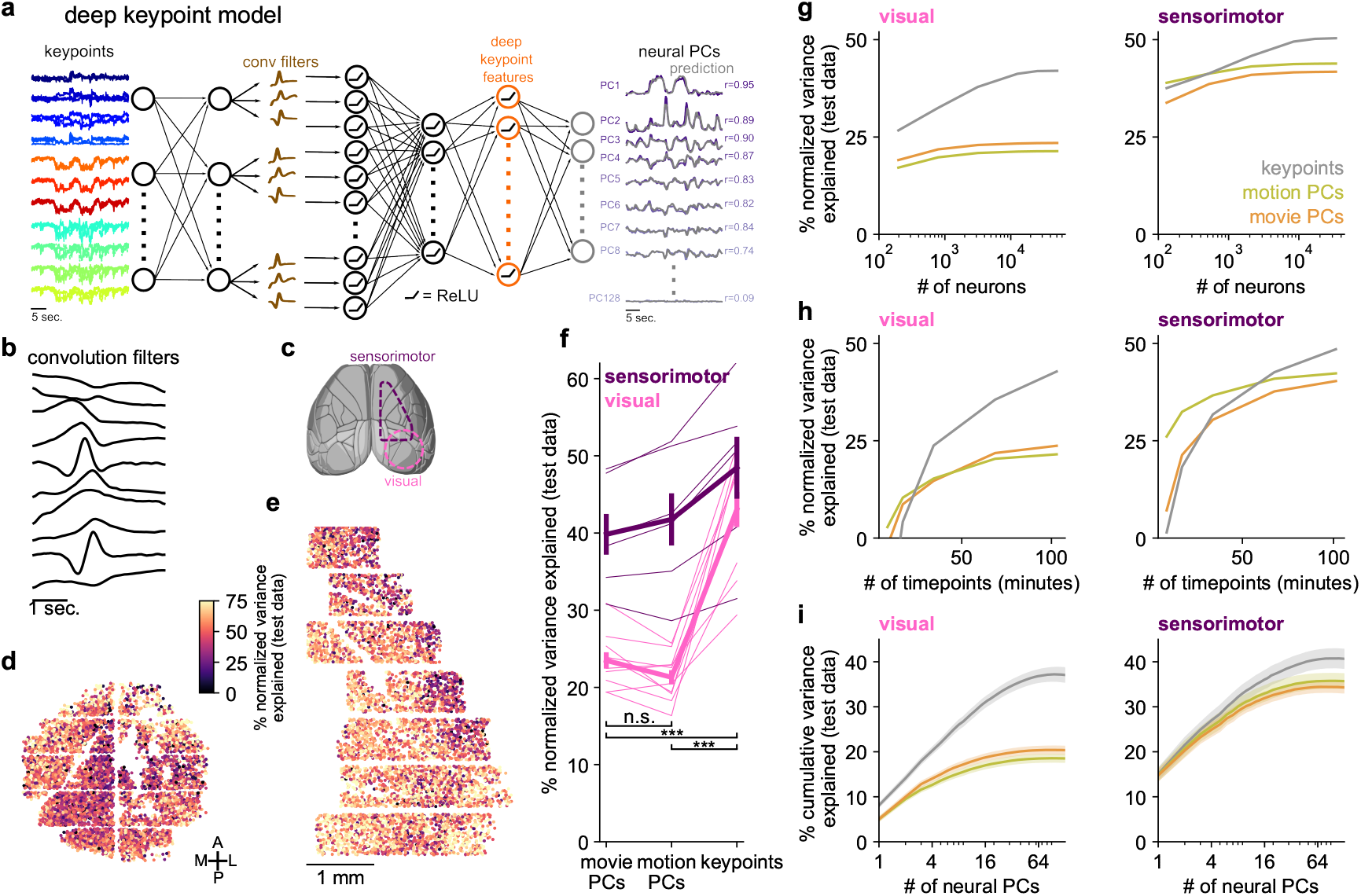
High-accuracy prediction of neural activity using keypoints. **a**, Architecture of 5-layer neural network for predicting neural activity. **b**, The resulting temporal convolution filters in layer 2 of the model for an example recording. **c**, Neural recording locations overlaid on the atlas from the Allen Institute for Brain Science (http://atlas.brain-map.org/). **d**, Neurons from an example recording in visual cortex colored by the percentage of normalized variance explained by the deep keypoint model on test data. **e**, Same as **d** for a recording in sensorimotor cortex. **f**, Percentage of normalized variance explained by movie PCs, motion PCs and keypoints, averaged across neurons for each recording. Thick lines denote average across recordings, error bars represent s.e.m., n=16 recordings. Two-sided Wilcoxon signed-rank test, *** denotes *p <* 0.001. **g**, Same as **f** as a function of the number of neurons, averaged across recordings from visual (left) and sensorimotor (right) areas. **h**, Same as **g**, for the number of timepoints in the training data. **i**, Cumulative variance explained across neural PCs from the keypoints, movie PC and motion PC predictions. Mean computed across n=16 recordings in 12 mice, error bars represent s.e.m.

We found that neurons across visual and sensorimotor cortices were well explained by behavior, with an average normalized variance explained of 43.0% and 48.5% respectively (Figure 3d,e). Next, we compared the neural prediction from keypoints to the neural prediction using our previous method based on movie and motion principal components (PCs) based on the raw mouse face videos. This method reduced the dimensionality of the movies from ∼100,000 pixels to 500 PC dimensions for neural prediction. The variance explained from the keypoints prediction was 92.2% higher than the variance explained from the movie and motion PCs in the visual recordings, and 18.8% higher in the sensorimotor recordings (Figure 3f). We also asked how the neural prediction varies with the number of neurons and timepoints available for the model fitting (Figure 3g,h). We found that explained variance saturated at around 10,000 neurons, but did not saturate even for our longest recordings of 2h. Thus, large-scale and longer recordings are necessary to fit good encoding models.

Finally, we noticed that the benefits of predicting from the keypoints were smaller for the sensorimotor recordings compared to the visual ones, and in fact the cells recorded most posterior in visual cortex had the largest improvement in prediction from keypoints, compared to anterior cells in higher order visual areas (Figure S4). We investigated the difference between visual and sensorimotor areas further by looking at variance explained individually per principal component (Figure 3i). First, we noticed that the benefit of the keypoints was primarily in explaining the higher neural PCs, both in the visual and sensorimotor areas. Second, we noticed that the first few PCs of the sensorimotor recordings accounted for a larger fraction of the total explained variance. For example, the first PC explained on average 15.1% of the variance in the sensorimotor recordings, but only 8.1% of the variance in the visual recordings. Thus, the additional benefit of the deep keypoint model in predicting visual cortex activity may be due to the higher-dimensional representations of behavior in this brain area.

The deep keypoint model preserved the finer-structure of neural activity, capturing small events across small groups of neurons better than the movie PC prediction (Figure 4a, Figure S5). We investigated the spatial distribution of these subgroups of neurons by clustering the neural activity into 100 clusters using k-means (Figure 4b,c). Some clusters were spatially spread throughout the recording area, while others were more localized (Figure 4c, Figure S6). We defined a spatial locality index for each cluster (see Methods). In general, the clusters best predicted by the behavior had the lowest locality index (Figure 4d,e). Thus, behaviorally-driven clusters are more spatially distributed across cortex, consistent with the hypothesis that many of these behavioral signals are broadcast across the brain.

**Figure 4:**
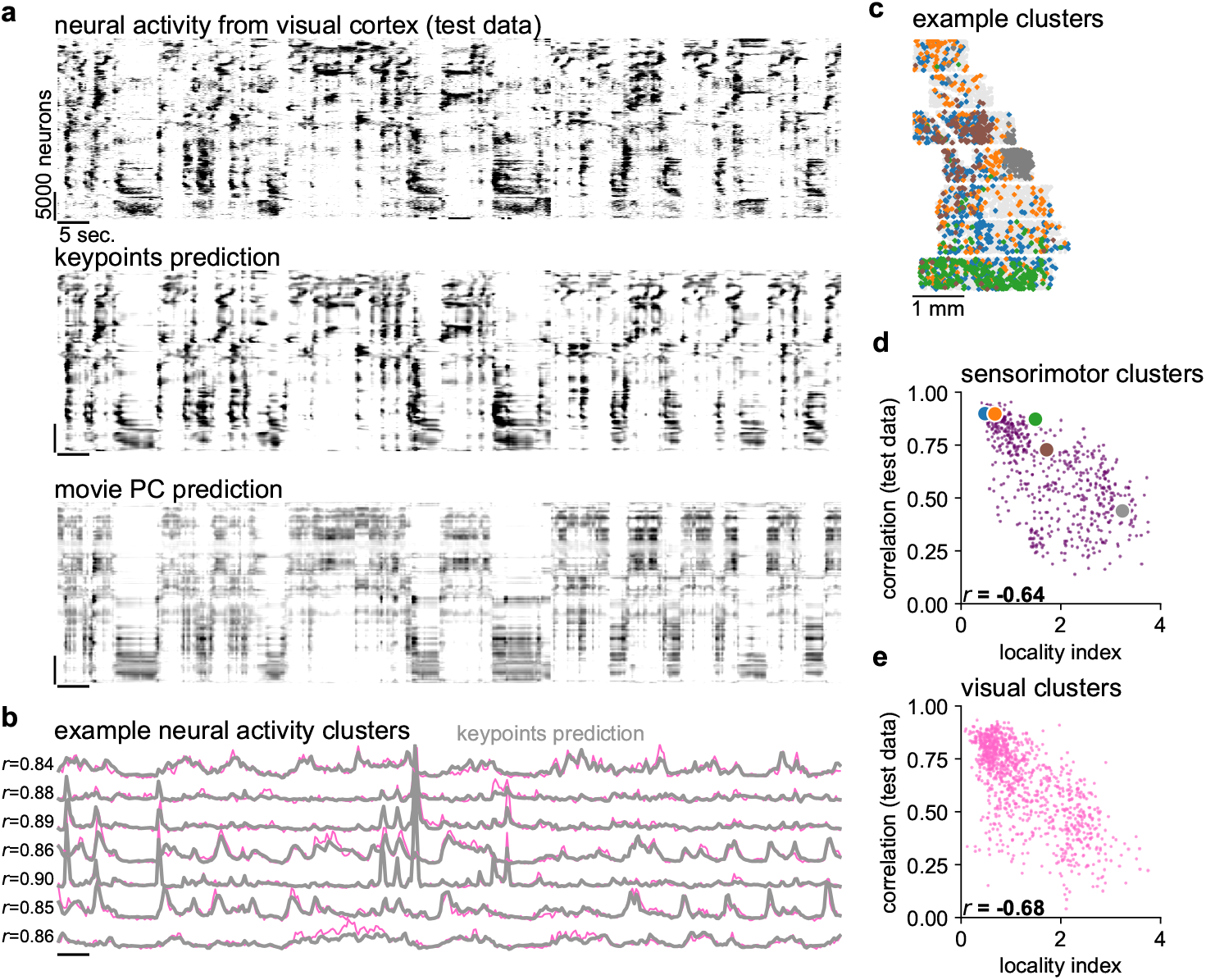
The deep keypoint model predicts fine features of neural activity. **a**, Top: Neurons from a visual (posterior, dorsal cortex) recording during spontaneous behavior. Each row represents averaged activity of 25 neurons, sorted using Rastermap [37]. The time period is during a held-out test period. Middle: Predicted neural activity from the deep keypoint model in Figure 3a. Bottom: Predicted neural activity using the movie PCs. **b**, Example neural activity clusters, same time period as **a. c**, Spatial locations of neurons from 5 example clusters from a recording in sensorimotor cortex. **d**, The locality index of each sensorimotor cluster across recordings, defined by the KL divergence between its spatial distribution and the distribution of all the neurons, plotted against the correlation of the cluster activity with its prediction from the deep keypoint model. The colored circles correspond to the clusters in **d. e**, Same as **d**, for visual clusters.

### State dynamics extracted from deep keypoint features

The last hidden layer in the deep keypoint model, the “deep keypoint features”, contain a representation of behavior that is not directly available in the raw keypoints. To understand the nature of these representations, we characterized their dynamical properties using Hidden Markov Models (HMM). Various types of HMMs have been previously fit to raw behavioral data, often from freely-moving animals [21, 39–42]. Here we chose to use discrete HMMs, which can model the data as a succession of discrete states [43, 44]. Transitions between states are probabilistic with probabilities defined by the transition matrix. In addition, each state is assigned a fixed “emission” pattern of activations across all features. The transition matrix and emission patterns are parameters that are fit to each session individually.

We start by visualizing HMM models that were fit to an individual session using 50 states. The 256 deep keypoint features from one session were first sorted using one-dimensional t-SNE, such that features with similar activation patterns are near each other in the plot (Figure 5a) [38]. The most probable states can then be inferred (Figure 5b), and their emission patterns can be used to reconstruct the original data matrix (Figure 5c). The reconstruction assures us that the HMM captures a majority of the data variance. We also visualized instances of the same state, and observed they were consistent and in some cases human-interpretable (Supplementary Videos 3). The HMM states were also separately sorted using

**Figure 5:**
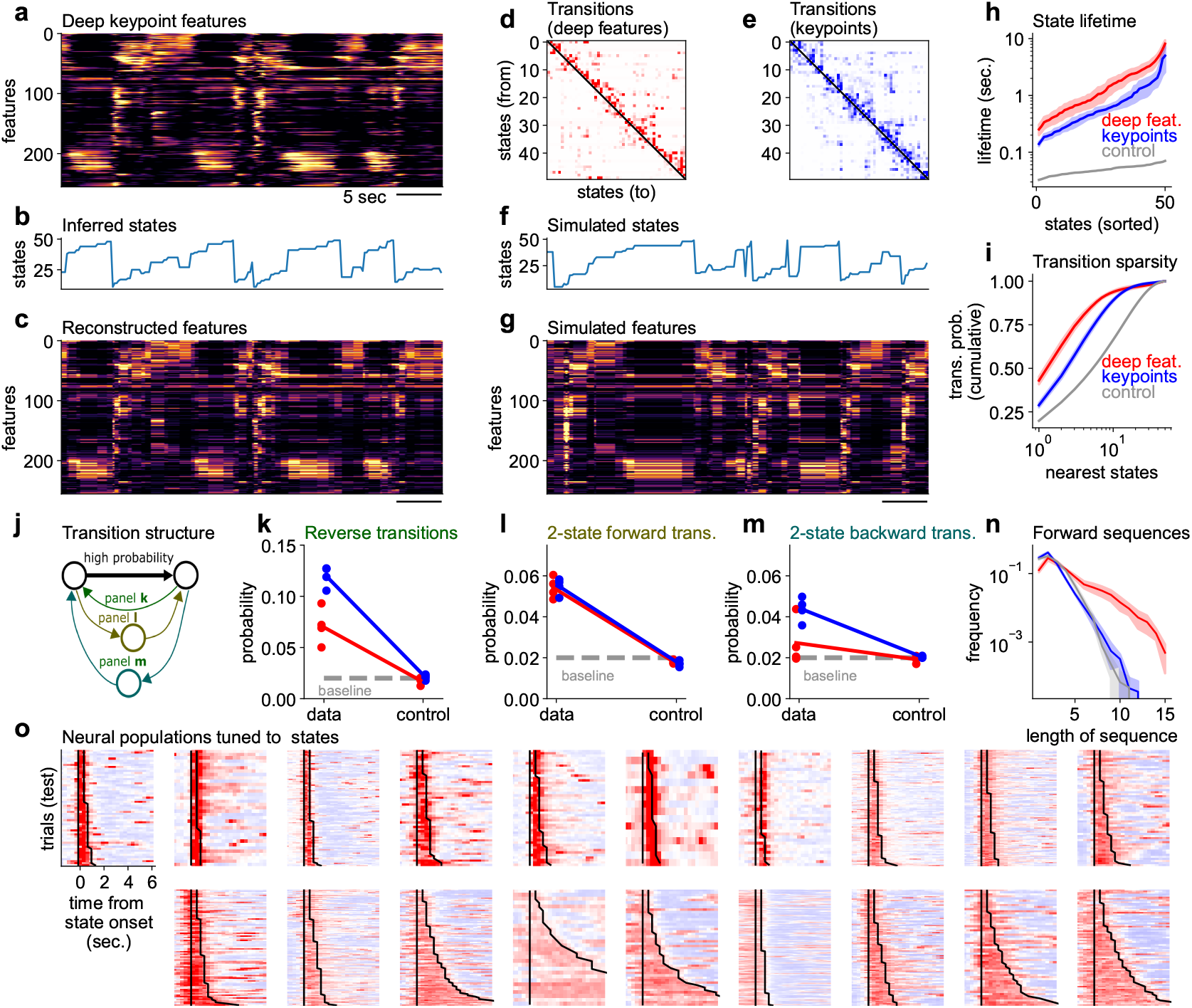
The deep keypoint features have highly-structured dynamics. **a**, Example dynamics of deep keypoint features computed by the neural network in figure 3**a**. The features have been sorted along the y-axis using a one-dimensional t-SNE embedding [38]. **b**, Inferred states using a Hidden Markov Model (HMM). **c**, Reconstructed features using the inferred states. **d**, State transition matrix of the HMM. Self-transitions were set to 0 and the rows were renormalized to 1. States have been sorted to maximize the sum of transition probabilities above the diagonal, using the Rastermap algorithm [37]. **e**, Same as **d** for HMMs fit directly to the keypoint data. **f**, Simulated states using the HMM fit to the deep keypoint features. **g**, Simulated features from the HMM. **h**, Distribution of inferred state lifetimes using the self-transition probabilities of the HMM. See Methods for description of controls for all panels. **i**, Probability of transitions to *N*-nearest states as a function of *N*. The average is taken over all initial states, and across mice. **j**, Schematic for panels **k**-**m**. For each pair of states with a high transition probability, certain other transition probabilities are reported. **k**, Average probability of reverse transitions. Baseline is computed as the average transition probability across all state transitions. **l**, Probability of two-state transitions. **m**, Probability of two-state backward transitions. **n**, Distribution of “forward” sequence lengths, where the forward direction is defined as higher indices in the Rastermap sorting of states from **d. o**, Neural populations tuned to 19 selected states (out of 50 total). Neurons were selected on train trials, and their average on test trials is shown. Vertical lines indicate trial onsets, while the second jagged line indicates trial offsets.

Rastermap [37], so that forward transitions – from a lower to a higher state in the sorting – are maximized in the sorting (Figure 5d). Due to this sorting, state dynamics appear to be arranged in ordered, increasing sequences (Figure 5b). This asymmetry in state transitions was not apparent at the level of the keypoints themselves (Figure 5e). Despite being sorted with the same Rastermap algorithm, states inferred directly from keypoints had relatively symmetric transition probabilities. To further validate the quality of the HMM, we used it to generate new synthetic data (Figure 5f,g). Samples from the model had the same overall appearance as the original data. Thus, transition probabilities captured in the HMM can generate the same kind of behavioral sequences as are present in the data itself.

Next, we quantified some of the HMM properties directly. The duration of a state in the model is given by the self-transition probability (which was left out from the visualizations in Figure 5d,e). Self-transitions *p* near 1, imply a long-lasting state, with an exponential distribution of state durations. The mean of this distribution is defined as the “state lifetime”, and can be easily computed as − log(1 − *p*). The distribution of state lifetimes was broad (Figure 5h), with lifetimes ranging from 0.2 to 10 seconds. The model fit to behavioral states had longer lifetimes than the model fit directly to the keypoints, and both had much longer lifetimes than a control model fit to temporally-shuffled data. For the rest of our analyses, we will ignore self-transition probabilities, and focus on the transition probabilities between states. Operationally, we set self-transitions to 0 in the transition matrix, and normalize the outgoing transitions to a sum of 1 (like in Figure 5d,e).

Another property of the HMM is the sparsity of transitions between states. It is apparent in Figure 5d that the transition matrix is quite sparse, with most values near-zero and a few large values. In other words, HMM states tend to transition to only a few other potential states. To quantify this property, we computed for each state the probability to transition to its *N* nearest states, where near states are defined as the ones with the highest probability of transition. As *N* increases, the summed transition probability approaches its maximum of 1 quickly for small *N* and much more slowly after. This shows that the HMM has a sparse structure, dominated by a few large transitions. In contrast, models inferred from the keypoints had more dense transitions (approached 1 more slowly). Both types of models had sparser transitions than the control model which was fit to appropriately-shuffled data (see Methods).

To quantify the asymmetry of the HMM transitions, we performed a series of analyses directly on the transition matrix (Figure 5j). For each pair of states with high transition probability, we ask how likely other transitions are. We analyzed reverse transitions (Figure 5k), two-step forward transitions (Figure 5l) and two-step backward transitions (Figure 5m). We found that these types of transitions were generally more likely than chance. However, reverse transitions were less likely in the deep feature HMM compared to the keypoint HMM, corresponding to the more asymmetrical nature of the former model (Figure 5d vs Figure 5e). While two-step forward transitions were matched between the two models, the two-step backward transitions were at baseline levels for the deep feature HMM, but not for the keypoint HMM. The net effect of the asymmetry in state transitions was that the deep feature HMM produced longer forward sequences of states. We quantified this property from the inferred states, measuring the length of all increasing state sequences (Figure 5n).The distribution of forward sequence lengths was much more long-tailed for the deep feature HMM, compared to controls and to the keypoint HMM (Figure 5n).

Finally, we visualized the activity of the neural populations tuned to different HMM states. We define a “trial” as uninterrupted timepoints of the same state, and the response of a neuron on that trial as its average activity over those timepoints. We then use training trials to select the neurons with the highest activity on each state. For each state, we thus obtain a neural population highly-selective to that state (Figure 5o). We observed populations with either brief or long-lasting activity, which mirrors the diversity of behavioral state durations. Other aspects of these neural populations could be investigated further, for example by engaging these neural populations in a behavioral task. However, that is beyond the scope of the present work.

## Discussion

Here we described Facemap, a framework that relates orofacial tracking to neural activity using new modeling tools. The framework is composed of two parts: 1) an orofacial keypoint tracker for extracting eye, whisker, nose and mouth movements, and 2) a neural network encoding model that extracts spatio-temporal features of behavior which are most related to the neural activity. We have shown that the orofacial tracker is highly accurate, while being significantly faster than other keypoint tracking approaches, and we showed that it can be easily trained on new orofacial videos from other experimental setups than our own. These keypoints capture the important aspects of the behavior with many fewer variables (22) than the number of pixels in a frame (∼ 100,000). Despite this dramatic dimensionality reduction, the keypoints contain enough information about the behavior to predict neural activity very accurately. Especially dramatic is the performance of the deep keypoint model in visual cortex, where it can predict almost twice as much variance compared to previous approaches.

We used the new Facemap framework to make a few initial observations. We found that the eye keypoints had predictable dynamics on much longer timescales (10 sec) compared to the dynamics of the nose keypoints (1 sec), while the whisker dynamics were somewhere in-between. We also found that visual cortex contained more dimensions of behavior compared to sensorimotor cortex, a surprising result that merits further investigation. Furthermore, the higher dimensions in visual cortex could not be predicted using our older,linear methods (movie and motion PCs), but instead required the nonlinear deep key-point model. This was not true in the sensorimotor areas, where the linear models performed almost as well as the new nonlinear model. Across both visual and sensorimotor areas, clusters that were spread out over the brain were the ones best predicted from behavior. Finally, we found that the deep keypoint features extracted by the model contained a much more orderly representation of behavior compared to the raw keypoints. Using an HMM, we found that the deep keypoint features were organized into relatively longer-lasting states, from less than a second to several seconds, which transitioned into other states in a predictable manner, forming sequences of states that repeated many times over the course of a session.

These initial analyses are just the start of using Facemap to extract insights about neural activity patterns and about the structure of behavior itself. We developed the method along with an user-friendly graphical user interface so that others can easily adapt it to their own data, and use it flexibly in their own studies. For example, many labs already have video cameras capturing the face of the mouse, and could therefore perform orofacial tracking during such experiments [45, 46]. Further, with head-mounted cameras [47], orofacial tracking could be incorporated into freely-moving behavioral contexts, to enable observation of the fine movements that rodents make as they explore their environment or engage in social interactions [39, 48–52]. We believe Facemap is one of the important steps towards unlocking the fundamental mystery of brainwide neural activity – what is its function, where is it coming from – and we look forward to seeing it used to make progress on these questions.

## Acknowledgments

This research was funded by the Howard Hughes Medical Institute at the Janelia Research Campus. We thank the Vivarium staff for animal husbandry, Sarah Lindo and Sal DiLisio for surgery support, Michalis Michaelos for help with recordings, Jon Arnolds for designing headbars and coverslips and Dan Flickinger for microscopy support. We thank Alice Robie, Mayank Kabra, and Kristin Branson for advice on keypoint tracking. We also thank the contributors of the mouse face videos from other labs: Adrian Hoffman in the Helmchen lab, Fanny Cazettes in the Mainen lab, Byron Price in the Gavornik lab, Aeron Laffere in the Lak lab, and Malcolm Campbell in the Uchida lab.

## Author contributions

A.S., C.S., and M.P. designed the study. A.S., C.S., M.P., and R.T. performed data analysis. L.Z., M.P., and W.L. performed data collection. A.S., C.S., and M.P. wrote the manuscript, with input from all authors.

## Code availability

Facemap was used to perform all analyses in the paper, the code and GUI are available at https://www.github.com/mouseland/facemap. Scripts for running the analyses in the paper are available at https://www.github.com/mouseland/facemap/paper/.

## Data availability

All data will be made available upon publication of the study.

## Methods

The Facemap code library is implemented in Python 3 [53], using pytorch, numpy, scipy, tqdm, numba, opencv, and pandas [34, 54–58]. The graphical user interface additionally uses PyQt and pyqtgraph [59, 60]. The figures were made using matplotlib and jupyter-notebook [61, 62].

### Data acquisition

#### Animals

All experimental procedures were conducted according to IACUC. We performed 16 recordings in 12 mice bred to express GCaMP6s in excitatory neurons: TetO-GCaMP6s x Emx1-IRES-Cre mice (available as RRID:IMSR JAX:024742 and RRID:IMSR JAX:005628). These mice were male and female, and ranged from 2 to 12 months of age. Mice were housed in reverse light cycle, and were pair-housed with their siblings before and after surgery. Due to the stability of the cranial window surgery, we often use the same mice for multiple experiments in the lab: 5 of the 7 visual area mice were used in a previous study [63], and the other two visual mice and all of the sensorimotor mice were trained on behavioral tasks after the recordings.

#### Surgical procedures

Surgeries were performed in adult mice (P35–P125) following procedures outlined in [63]. In brief, mice were anesthetized with Isoflurane while a craniotomy was performed. Marcaine (no more than 8 mg/kg) was injected subcutaneously beneath the incision area, and warmed fluids + 5% dextrose and Buprenorphine 0.1 mg/kg (systemic analgesic) were administered subcutaneously along with Dexamethasone 2 mg/kg via intramuscular route. For the visual cortical windows, measurements were taken to determine bregma-lambda distance and location of a 4 mm circular window over V1 Cortex, as far lateral and caudal as possible without compromising the stability of the implant. A 4+5 mm double window was placed into the craniotomy so that the 4mm window replaced the previously removed bone piece and the 5mm window lay over the edge of the bone. The sensorimotor window was also a double window and it was placed as medial and frontal as possible. The outer window was 7mm by 4.5mm and the inner window was around 1mm smaller in all dimensions. After surgery, Ketoprofen 5mg/kg was administered subcutaneously and the animal allowed to recover on heat. The mice were monitored for pain or distress and Ketoprofen 5mg/kg was administered for 2 days following surgery.

#### Videography

The camera setup was similar to the setup in [14]. Infrared LEDs (850nm) were pointed at the face and body of the mouse to enable infrared video acquisition in darkness. The videos were acquired at 50Hz using FLIR cameras with a zoom lens and an infrared filter (850nm, 50nm cutoff). The wavelength of 850nm was chosen to avoid the 970nm wavelength of the 2-photon laser, while remaining outside the visual detection range of the mice.

#### Imaging acquisition

We used a custom-built 2-photon mesoscope [64] to record neural activity, and ScanImage [65] for data acquisition. We used a custom online Z-correction module (now in ScanImage), to correct for Z and XY drift online during the recording. As described in [63], we used an upgrade of the mesoscope that allowed us to approximately double the number of recorded neurons using temporal multiplexing [66].

The mice were free to run on an air-floating ball. Mice were acclimatized to running on the ball for several sessions before imaging. On the first day of recording, the field of view was selected such that large numbers of neurons could be observed, with clear calcium transients.

#### Processing of calcium imaging data

Calcium imaging data was processed using the Suite2p toolbox [67], available at www.github.com/MouseLand/suite2p, which relies on the packages numpy, scipy, numba, scanimage-tiff-reader, paramiko, and scikit-learn [68–72]. Suite2p performs motion correction, ROI detection, cell classification, neuropil correction, and spike deconvolution as described elsewhere [14]. For non-negative deconvolution, we used a timescale of decay of 1.25 seconds [73, 74]. We obtained 50,614 ± 13,919 (s.d., n=10 recordings) neurons in the visual area recordings, and 33,686 ± 4465 neurons (n=6 recordings) in the sensorimotor area recordings.

### Facemap tracker network

#### Model architecture

The Facemap tracker network is a U-Net-style convolutional neural network consisting of downsampling and upsampling blocks with skip connections implemented in pytorch [34]. The model’s input is a grayscale 256 × 256*px* image, which is passed through a set of convolutional filters of different sizes shown in Figure 1b. The network has two sets of outputs: (i) heatmaps representing probability of a key-point in the pixel region, and (ii) location refinement maps representing the *x* and *y* offsets between the keypoint position in full-sized image and the down-sampled map, similar to [27, 33]. The downsampled (64 × 64*px*) heatmaps and location refinement maps are used to obtain the *x* and *y* coordinates of keypoints, example traces shown in Figure 1f. The

The tracker predicted 15 distinct keypoints in total for tracking mouse orofacial movements from different views (Figure 1a, Figure S1). The keypoints were used to track various movements of the eye (4), nose (5), whiskers (3), mouth (2) and an additional keypoint for the paw. The forepaw occasionally entered the view, such as during grooming, but we found this keypoint difficult to track and use in further analyses, so we did not consider it further. We also labeled a fifth nose keypoint (nosebridge, not shown in Figure 1a and Figure S1), but found that it was difficult to identify across different camera angles and therefore excluded it from the analyses in the paper. The videos taken during neural recordings were from the view in Figure 1c. In this view, the mouth keypoints were not visible, so those keypoints were not used in the model for neural prediction. Thus, we used 4 eye keypoints, 4 nose keypoints, and 3 whisker keypoints for neural prediction.

#### Training

The Facemap tracker was trained on 2400 images recorded from multiple mice and different camera views (Figure S1). Training images of size 256 × 256 pixels were labeled with all the keypoints, except when a bodypart was not visible in frame then no label was added. The model was trained for 36 epochs with the Adam optimizer using a batch size of 8 and weight decay of zero [75]. We used custom learning rate scheduler that used a fixed learning rate (*LR*) of 0.0004 for 30 epochs followed by 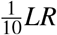 for the next 3 epochs and finally 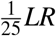 for the final 3 epochs. Each image was normalized such that 0.0 represented the 1^st^ percentile and 1.0 represented the 99^th^ percentile. Image augmentations performed during training were random crop, resize after padding to maintain aspect ratio, horizontal flip, and contrast augmentation.

#### Performance evaluation

Accuracy of the tracker was evaluated using the average pixel error for 100 test frames of size 256 × 256 pixels from a new mouse and different camera views. First, the Euclidean distance in pixels between the ground-truth labels and the predicted keypoints was computed. Next, the average error was computed as the average of the Euclidean distances across all frames (Figure S2a) and across all keypoints (Figure 1e).

Inference speed of the tracker was calculated to evaluate its utility for offline and online analysis. Hence, the inference speed calculation accounted for timing of various steps: (i) image pre-processing, (ii) forward pass through the model, and (iii) post-processing steps. All inference speeds are reported for a sample image of size 256 × 256 pixels passed through the network for 1024 repetitions and a total of 10 runs using various batch sizes on different GPUs (Table S2).

#### Filtering keypoint traces for neural prediction

Occasionally keypoints are occluded, such as during grooming. Therefore, like DeepLabCut, we found the timepoints when the tracker network confidence was low, and replaced those timepoints in the keypoint traces by a median filtered value. The network confidence, or likelihood, is defined as the value of the peak of the heatmap output. The likelihood traces for each keypoint were baseline filtered in time with a gaussian filter of standard deviation 4 seconds, then the threshold of the likelihood was defined as -8 times the standard deviation of the baselined likelihood, and any values below this threshold were considered outliers. This identified on average 0.19% of timepoints across all keypoint traces as outliers.

After excluding outliers based on likelihood, we also directly identified outlier timepoints using the keypoint traces, by detecting large movements or deviations from baseline. If a keypoint moved more than 25 pixels from the previous timepoint to the current time-point, then the current timepoint was considered an outlier. Also if the keypoint trace on the current time-point exceeded its median filtered (window=1 second) value by more than 25 pixels, then the current time-point was considered an outlier. This identified on average an additional 0.066% timepoints across all key-point traces as outliers.

To obtain values for the outlier timepoints, we median-filtered the keypoint traces with a window of 300 ms, excluding the outlier timepoints. Linear interpolation from the median-filtered traces was then used to fill in the values at the outlier timepoints.

### Pose estimation model comparisons

We compared the performance of the Facemap tracker to other state-of-the-art tools used for pose estimation, including SLEAP [31] and DeepLabCut [27]. The models were trained on the same training set used for Facemap. In addition, the same protocol for speed benchmarking was used to obtain the inference speed of the other models.

#### DeepLabCut models training

DeepLabCut’s models used for comparison included two different architectures: ResNet50 (default model) and Mobilenet v2 0.35 (fastest model). Augmentations used during training were scaling, rotation and contrast augmentation, similar to training of the Facemap tracker. A hyperparameter search was performed to find optimal training parameters for each model using different batch sizes (1, 2, 4, 8) and learning rates (0.0001, 0.001, 0.01). Models with the lowest average test error for each architecture were compared to Facemap in Figure 1e and Figure S2.

Inference speeds for DeepLabCut’s models were obtained using similar approach as Facemap tracker. We timed DeepLabCut’s *getposeNP* function for 1024 repetitions for a total of 10 runs for different batch sizes and GPUs. The *getposeNP* function timing included a forward pass through the network and post-processing steps to obtain keypoints locations from the heatmaps and location refinement maps.

#### SLEAP models training

The default U-Net backbone was used for SLEAP’s models, which included two different values of initial number of filters: (i) *c* = 16 (default) and (ii) *c* = 32 to vary the network size and potentially improve accuracy. A hyperparameter search over different learning rates (0.0001, 0.001, 0.01), batch sizes (1, 2, 4, 8) and number of epochs (100, 150) was performed to find the best model for each U-Net configuration. Furthermore, early stopping by stopping training on plateau was used for half of the models to prevent overfitting. Default augmentation settings were used for most models and mirroring (horizontal flip) was added to some models to match training of the other networks used for comparison. Similar to DeepLabCut, the best models were selected based on the lowest average test error for the default and *c* = 32 models, and used in Figure 1e and Figure S2.

Inference speed for SLEAP’s models was calculated by timing their *predict on batch* function. The U-Net models with different number of initial filters were run for 1024 repetitions for a total of 10 runs using different batch sizes of our sample image input.

### Facemap tracker refinement

We developed a method for fine-tuning the Facemap tracker for new data which differed from our training data. Facemap tracker’s base model is defined as the network trained on our dataset (Figure 1). We extracted frames from videos contributed by five other labs to use as training data for fine-tuning the base model specifically to each lab’s video. We executed the following steps for each lab’s video. First, the base model was used to generate predictions for 50 random frames. Keypoints on the 50 training frames were refined to correct keypoints with large deviations from their defined bodyparts or remove keypoints not in view. Next, the base model was fine-tuned with varying numbers of training frames ranging from 1-50. The network was trained for 36 epochs with an initial learning rate of 0.0001 with annealing as described earlier and weight decay of 0.001. Additionally, we trained a model from scratch, i.e. a network initialized with random weights, using 10 training frames for comparison. To compute the errors for the base model, the finetuned model, and the scratch model, we used 50 test frames and labeled them from scratch to use as a test set. We then computed the average error in pixels from the test set labels to the model predictions (Figure 2b). The models trained from scratch with 10 frames had an average error of 3.76 ± 0.39 pixels across labs, compared to 2.43 ± 0.24 pixels for the base model fine-tuned with 10 frames. Predictions from the base, scratch, and fine-tuned models for a random section of the video are shown in Supplementary Videos 2 for each lab. The workflow used for the analysis was integrated into the GUI so users can easily fine-tune the Facemap tracker with video recordings which differ from our training data (Figure 2c).

### Behavior to neural prediction

The activity of each neuron was z-scored: the activity was subtracted by the mean and divided by the standard deviation. In order to predict the neural activity from behavior, we reduced the dimensionality of the z-scored activity using principal component analysis (PCA) to obtain 128 neural PCs (*Y*). If fewer than 200 neurons were predicted, then we directly predicted the neurons rather than using the PCs.

The neural activity was split into 10 segments in time. The first 75% of each segment was assigned to the training set, and then after 3 seconds which were excluded, the remaining part of the segment was assigned to the test set. The training and test set were made to consist of continuous segments to avoid contamination of the test set with the train set due to the autocorrelation timescale of behavior, with lengths on average of 10 minutes and 3.5 minutes respectively.

We quantified the performance of a neural prediction model using the variance explained. The single neuron variance explained for a neural trace for neuron *i* (**s**_*i*_) is defined as

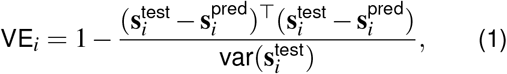

which is the standard definition for variance explained.

#### Peer prediction analysis

Neurons have independent noise that models cannot explain. Therefore, an upper bound for the variance that a model can explain is lower than the total variance of neural activity. To estimate the amount of this explainable variance in the neural recordings, we used the “peer prediction” method [14, 76, 77]. Peer prediction analysis predicts each neuron from the other simultaneously recorded cells (the neuron’s “peers”). The amount of variance that the peer prediction model is an estimate of the repeatable shared variance across neurons, we term this variance the explainable variance.

To compute peer prediction, we split the population into two spatially segregated populations, dividing the field of view into non-overlapping strips of width 200 *μ*m and assigning the neurons in the even strips to one group, and the neurons in the odd strips to the other group, regardless of the neuron’s depth. Next we computed the top 128 principal components of each population and predicted one population’s PCs from the other population’s PCs using reduced rank regression fit to training data with λ=1e-1 and rank=127. The variance explained by this model on test data (Equation 1) is termed the *explainable variance* for each neuron. The average explainable variance was 9.4% in the visual recordings and 11.1% in the sensorimotor recordings at the recording frame rate 3 Hz.

#### Prediction performance quantification

We computed the variance explained for a given behavioral prediction model for each neuron on test data (Equation 1). The average single neuron variance explained by the deep keypoint model was 4.1% in the visual areas and 5.2% in the sensorimotor areas at 3 Hz. We then normalized the variance explained by the upper bound on its variance explained, the explainable variance, as computed from peer prediction. We quantified the normalized variance explained on a per neuron basis in Figure 3d,e, taking the variance explained for each neuron and dividing it by its explainable variance, and visualizing only the neurons with an explainable variance greater than 1e-3. For population-level variance explained quantification, the normalized variance explained was defined as the mean variance explained across all neurons divided by the mean explainable variance across all neurons (Figure 3f-i, Figure S3, Figure S4).

We also computed the cumulative variance explained across neural PCs (**y**_*i*_), defined as

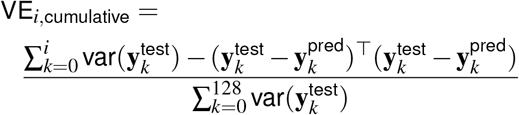

in Figure 3i. This quantity allows the estimation of dimensionality of the behavioral prediction.

#### Neural prediction using PCs of videos

The mouse videos were reduced in dimensionality using singular value decomposition (SVD) in blocks as described in [14]. The movie PCs were computed from the raw movie frames, and the motion PCs were computed from the absolute value of the difference between frames. For each, the top 500 PCs were used. Because the neural activity was recorded at a lower frame rate, the behavioral PCs were smoothed with a Gaussian filter of width 100 ms and then resampled at the neural timescale. We subtracted each behavioral PC by its mean, and divided all PCs by the standard deviation of the top behavioral PC.

A linear model called reduced rank regression (RRR) was used to predict neural PCs (*Y*) from behavioral PCs (*X*). Reduced-rank regression is a form of regularized linear regression, with the prediction weights matrix restricted to a specific rank [78], reducing the number of parameters and making it more robust to overfitting. The RRR model is defined as:

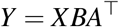

Like in ridge regression, the identity matrix times a λ constant can be added to the input covariance *X* for regularization. We set λ=1e-6. Training data was then used to fit the *A* and *B* coefficients in closed form, and a rank of 32 was used.

#### Neural prediction using keypoints

A multi-layer network model was fit to predict neural activity from corrected keypoints traces using pytorch [34] (Figure 3a). The deep keypoint model consisted of a core module and a readout module. The core module consisted of a fully-connected layer with the same dimensionality as the number of keypoints, a one-dimensional convolutional layer with 10 filters (temporal convolution), a ReLU non-linearity, two fullyconnected layers with ReLU non-linearities, the first with dimensionality 50 and the second with dimensionality 256. The 256-dimensional output of the core module is termed the “deep keypoint features” of the model. The readout module of the network was one fully-connected layer, with dimensionality of size 128 when predicting the neural PCs, or size equal to the number of neurons when predicting single neuron activity (when the number of neurons predicted was less than 200). The deep keypoint features, before entering the readout module, were subsampled at the time-points coincident with the neural activity frames, because the videos were recorded at 50Hz while the neural activity was recorded at 3Hz.

The deep keypoint model was fit on the training data using the optimizer AdamW with learning rate 1e-3, weight decay 1e-4, and 300 epochs [79], and the learning rate was annealed by a factor of ten at both epoch 200 and epoch 250. When fewer than 2000 neurons were fit, the learning rate and weight decay were reduced by a factor of ten to reduce over-fitting. When fewer than 1 hour of training timepoints were used, the learning rate and weight decay were reduced by a factor of two and the number of epochs reduced by 100 to reduce overfitting. Each training batch consisted of a single training segment, on average of length ten minutes, and there were ten batches per recording. The model was then applied to the test segments to compute variance explained.

We varied various parameters of the network to approximately determine the best network architecture for neural prediction Figure S3. We varied the number of units in the last layer of the core module, the “deep keypoint features”, from 1 to 1024 (Figure S3a), and the number of convolution filters (Figure S3e). We varied the number of fully-connected layers with ReLU non-linearities in the core module, each with dimensionality 50 other than the last layer which was fixed at 256 dimensions (Figure S3b). We also varied the number of fully-connected layers in the readout module, with each layer having 128 dimensions and a ReLU non-linearity, other than the last layer which had no output non-linearity (Figure S3c). Next, from the original architecture described above, we removed components, such as the first fully-connected layer and some of the ReLU non-linearities (Figure S3d).

#### Scaling of performance with neurons and timepoints

In Figure 3g-i, we quantified the prediction performance as a function of the number of neurons and timepoints. For this analysis we predicted using either a fraction of the neurons or a fraction of the training timepoints, while always keeping the test timepoints fixed. The variance explained was computed for each neuron, averaged across all neurons in the subset, and then normalized by the explainable variance averaged over the neurons in the subset.

### Neural activity clustering and sorting

We identified groups of coactive neurons usings scaled k-means clustering [67]. Compared to regular k-means, scaled k-means fits an additional variable λ_*i*_ for each neuron *i* such that

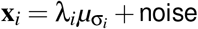

where **x**_*i*_ is the activity vector of neuron *i*, σ_*i*_ is the cluster assigned to neuron *i* and *μ*_*j*_ is the activity of cluster *j*. Like regular k-means, this model is optimized by iteratively assigning each neuron to the cluster which best explains its activity, and then re-estimating cluster means. We ran scaled k-means clustering with 100 clusters on z-scored neural activity. Example clusters shown in Figure 4c and Figure S6. The activity of the neurons in each cluster was averaged to obtain a cluster activity trace (Figure 4b). To obtain the cluster prediction from the deep keypoint model, we averaged the prediction of each neuron in the cluster (shown in gray in Figure 4b), and then correlated this prediction with the cluster activity trace to obtain an *r* value for each cluster.

To quantify how spread out each cluster is in the recording field of view, we computed a locality index for each cluster. We defined the locality index as the Kullback-Leibler (KL) divergence between the cluster’s discretized spatial distribution in the recording field of view and the discretized spatial distribution of all neurons, using a discretization of 200 *μ*m. We then correlated the locality index with the correlation of each cluster with its prediction (Figure 4d,e).

### Fitting a discrete Hidden Markov Model

We fit Hidden Markov Models (HMM) to the deep key-point features {*z*_*t*_}_*t*_, where *t* is a time-step for temporal features that were downsampled 10× from 50 Hz to 5 Hz [43]. We also fit the same models to the keypoint data. All fitting procedures were the same, except for the choice of the variance term, which depends on the number of features (30 for the 11 keypoints from the Facemap tracker and 256 for the deep keypoint features) in a way described below. The HMM state dynamics are given by:

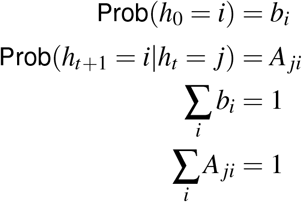

where *b*_*i*_ represents the probability of starting the Markov chain in state *i*, while *A*_*ji*_ represents the probability of transition from state *j* to state *i*. In all experiments we chose the number of states to be 50, and so similar results with fewer (10) or more (200) states. Since our goal is to understand the pattern of dynamics of the deep keypoint features, we did not attempt to infer the “optimal” number of states and do not believe the data lends itself easily to such an estimation.

In addition to state dynamics, an HMM has an “observation” or “emission” model, which declares the probability of observing some data sample *z*_*t*_ for each possible state *h*_*t*_ :

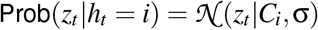

where *C*_*i*_, σ are the mean and standard deviation of the Gaussian observation model. This completes the model specification. We optimized this model in pytorch using an improved, non-standard optimization scheme, which routinely optimized the model better compared to alternative optimization methods such as expectation maximization.

Our optimization scheme consists of 1) optimizing the model log-likelihood directly as a function of its parameters using the automated differentiation from pytorch and 2) using initializations and reparametrizations of the HMM parameters that improve stability.

The log-likelihood of the HMM can be computed based on the forward pass of the “forward-backward” algorithm. Following the convention of [44], we define α(*h*_*t*_) = Prob(*z*_1_, *z*_2_, …, *z*_*t*_, *h*_*t*_). We can then define recursion equations for computing

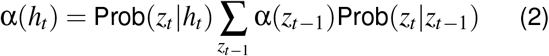

The full log-likelihood of the data can then be computed based on α(*h*_*T*_), where *T* is the last timepoint, by observing that

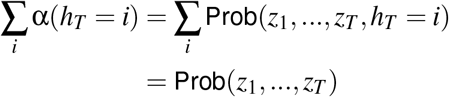

Since the dependence of α_*t*+1_ on α_*t*_ can be written in closed form, we can see that they are differentiable. After taking the logarithm and replacing the probabilities with the model equations, Equation 2 becomes:

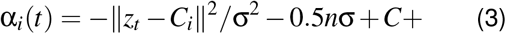

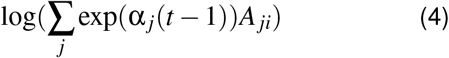

where α_*i*_(*t*) = log(α(*h*_*t*_ = *i*)), *C* is a constant and *n* is the number of dimensions of the data. This formulation allows us to use the automatic differentiation from pytorch to optimize the HMM model directly, without inferring states first like in the expectation maximization method. Additionally we note that we used the “logsumexp” function from pytorch to compute the second half of Equation 4, which has the advantage of being stable to exponentiation.

We re-parametrized the transition matrix *A* with a “log-transition” matrix *Q* by

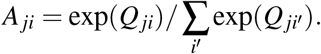

This has the advantage of removing the constraint of positivity of *A*_*ji*_ and the summing to 1 of the rows of *A*. We initialized the log-transition matrix with *Q*_*ii*_ = 3 and *Q*_*i j*_ = 0 when *i* ≠ *j*, and we initialized the parameters *C*_*i*_ of the observation model with random samples from the data. For setting σ, we made the choice of freezing it to a fixed value for each dataset. This was because of the dependence of the log-likelihood on the number of observation dimensions *n* in Equation 4. Since *n* is quite different between the keypoints and the deep keypoint features, the relative contribution of the observation term to the likelihood would be different if we set or learned σ to be the same in the two cases, potentially biasing the model to rely more or less on the internal hidden states *h*_*t*_. Instead, we fix σ^2^ to be proportional to the summed variance of *z*_*t*_, and we set it to 1 for the deep keypoint features, and 30*/*256 for the keypoints model. This ensures an approximately equal weighting of the observation term into the likelihood model. We note that the properties of the fitted HMM were not substantially different when σ^2^ was set to the same value for the keypoints and deep keypoint features, but the quality of the samples simulated from the HMM degraded if σ^2^ was too low.

### Properties of the discrete Hidden Markov Model

The inferred states were determined with the Viterbi algorithm, which finds the most likely hidden states. We simulated states by drawing initial states from the categorical distribution with parameters *b*_*i*_, and then running the forward dynamics and drawing states from the conditional distributions Prob(*h*_*t*+1_ = *i*| *h*_*t*_ = *j*) = *A*_*ji*_.

State lifetimes were defined as −log(1 −*A*_*ii*_), and they correspond to the mean durations of staying in state *i*. To compute transition sparsity and other metrics, we set self-transitions *A*_*ii*_ = 0 and renormalized the rows. Formally, we defined a transition matrix *B*_*ji*_ = *A*_*ji*_*/* ∑_*i*_′≠ *j A*_*ji*_′ when *j* ≠ *i* and *B*_*ii*_ = 0. This is the matrix shown in Figure 5de and using for the analyses in Figure 5i-n. The states were sorted using the Rastermap algorithm on the matrix *B* [37]. Specifically, this involves maximizing the similarity of the reordered transition matrix to the matrix given by *F*_*ji*_ = −log((*i* − *j*)^2^) when *j < i* and 0 otherwise. Thus, the model attempts to put the highest probabilities close to the diagonal, and specifically above the diagonal, since they don’t count if they are below the diagonal. For more details, see the rastermap repository at github.com/MouseLand/rastermap.

The transition sparsity was computed by sorting the rows of the matrix *B* in descending order, and computing a cumulative sum over each row. “Near” states were defined as the five states *i* with highest probability *A*_*ji*_ for a given *j*. Reverse transitions were computed for each state based on its near states. Similarly we computed the 2-state forward and backward transitions. Forward sequences were computed based on the most likely inferred states, by counting the number of increasing sequences of each length. Note this depends on the initial Rastermap sorting of states to define a meaningful order.

---

**Table S1:**
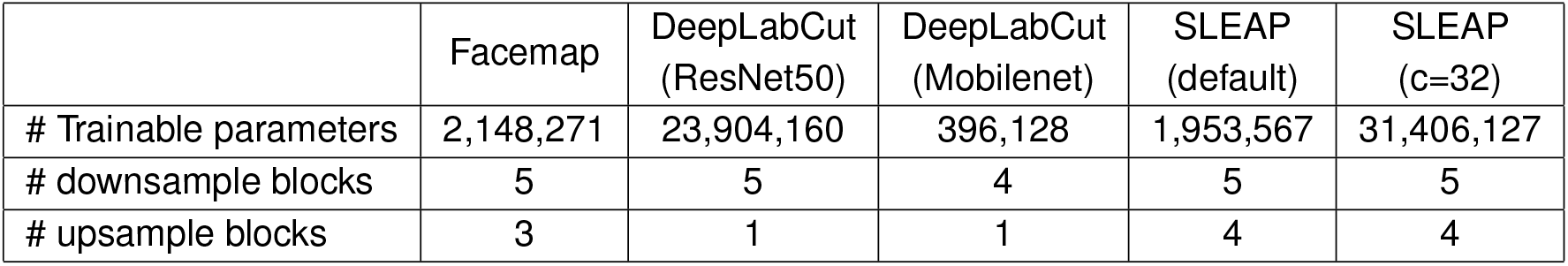
Network architecture details of the Facemap tracker, DeepLabCut and SLEAP models. Number of trainable parameters and the number of downsample and upsample blocks used for each model.

**Table S2:**
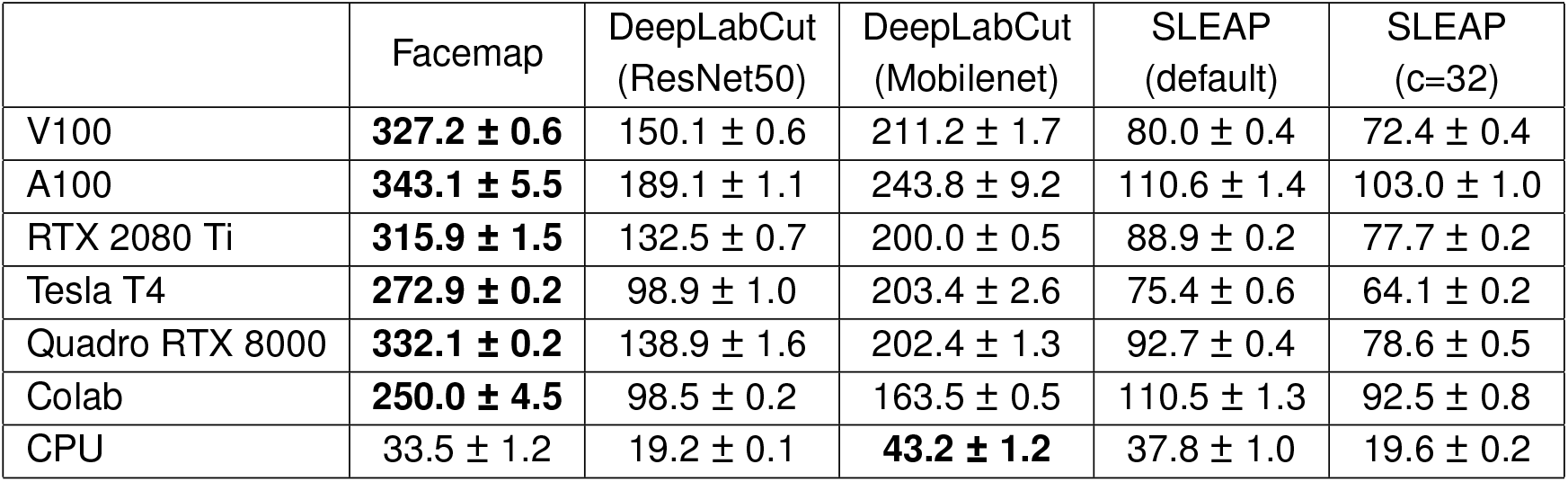
Inference speed of pose estimation models on different GPUs. A sample image of size 256 × 256 pixels was used as input for Facemap, DeepLabCut (ResNet50), DeepLabCut (Mobilenet), SLEAP (default) and SLEAP (c=32) models. The time taken for a forward pass through the network and post-processing steps was used to obtain inference speed for batch size of 1. A total of 1024 repetitions and 10 runs were used to obtain the average inference speed. The number of CPU cores/slots used for different GPUs were as follows: A100 (48 slots, 40GB/slot), V100 (48 slots, 30GB/slot), RTX 2080 Ti (40 slots, 18GB/slot), Tesla T4 (48, 15GB/slot) and Quadro RTX 8000 (40 slots, 18GB/slot). The CPU used was 3.0GHz Intel Cascade Lake(Gold 6248R) with 768GB and 8 slots used for timing on CPU only. Models were also run on Google Colab’s GPU.

**S1:**
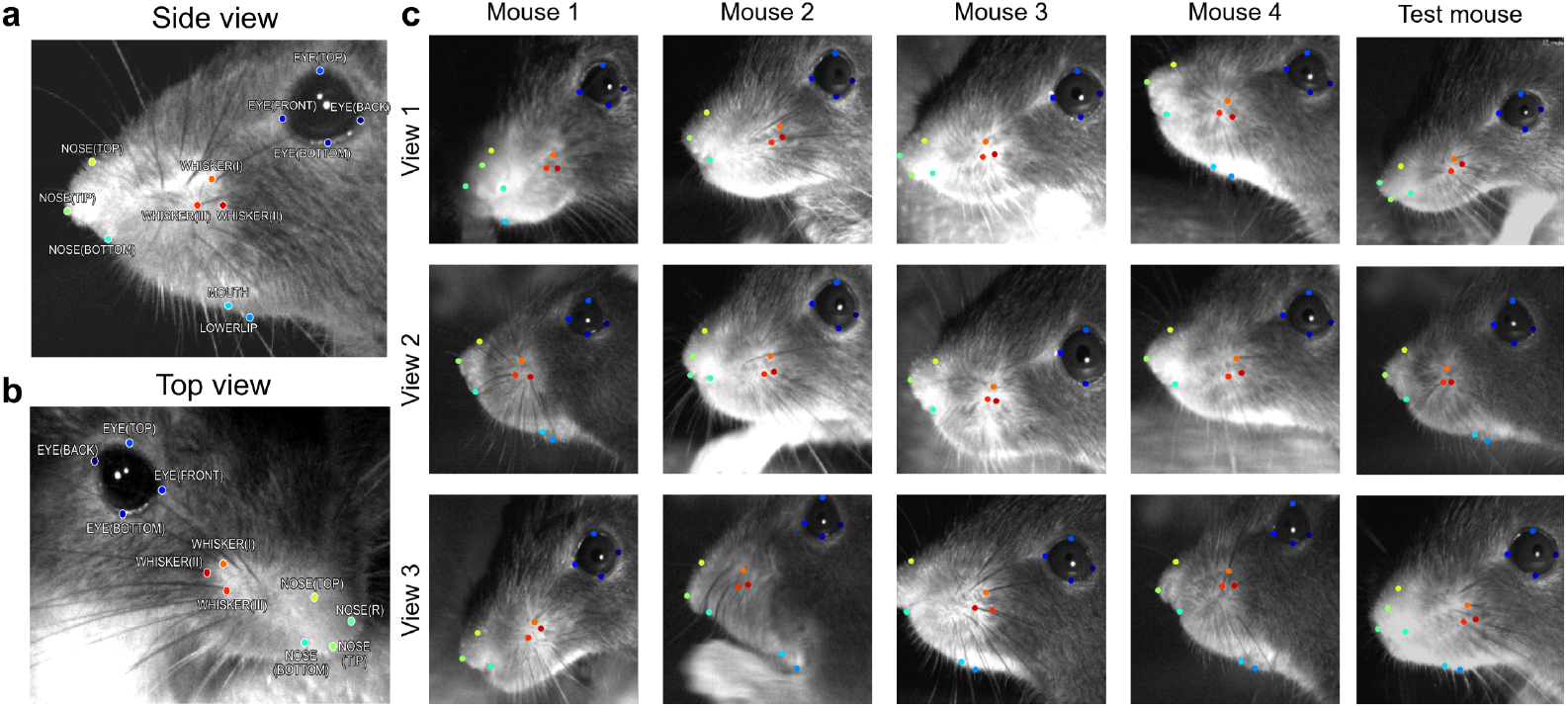
Mouse face keypoints from different camera views. Keypoints labels and sample images shown from training and test set. **a**, Side view recording of a mouse face showing eye, whiskers, nose and mouth keypoints. **b**, Top view recording of mouse face in **a** showing eye, whiskers and nose keypoints from a different view. **c**, Mouse face recordings from different camera views for training samples and test samples (last column). A total of 2400 training frames were used from mice shown in **c** and other mice (not shown), and 100 frames from different views of a new mouse used as the test set.

**S2:**
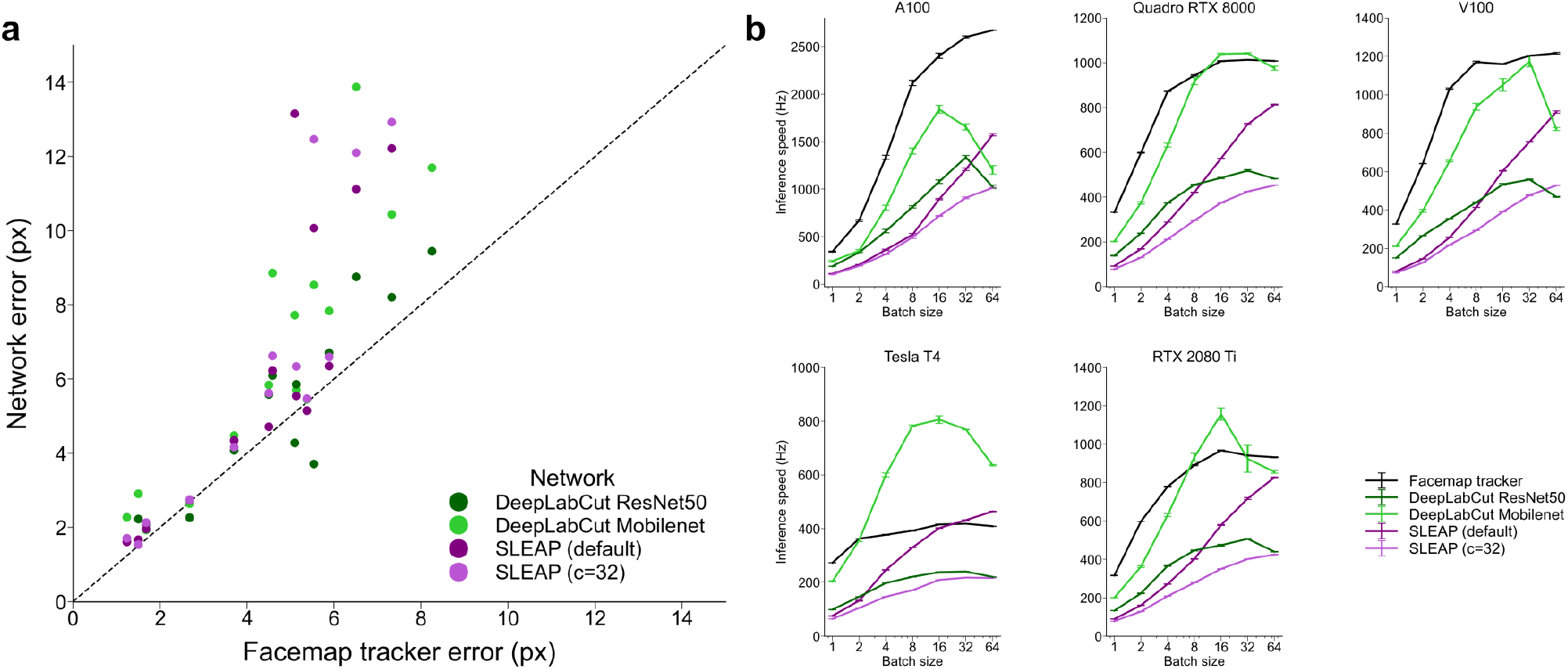
Per keypoint error and inference speed of networks using various batch sizes. **a**, Error for each keypoint, averaged across 100 test frames for each network plotted against the Facemap tracker errors. Points above the diagonal indicate keypoints for which Facemap outperformed the other networks. **b**, Inference speed of Facemap, DeepLabCut (ResNet50), DeepLabCut (Mobilenet), SLEAP (default) and SLEAP (c=32) models for a sample image of size 256 × 256 pixels on A100 (48 slots, 40GB/slot), V100 (48 slots, 30GB/slot), RTX 2080 Ti (40 slots, 18GB/slot), Tesla T4 (48, 15GB/slot) and Quadro RTX 8000 (40 slots, 18GB/slot). Inference speed averages shown for a total of 1024 repetitions across 10 runs.

**S3:**
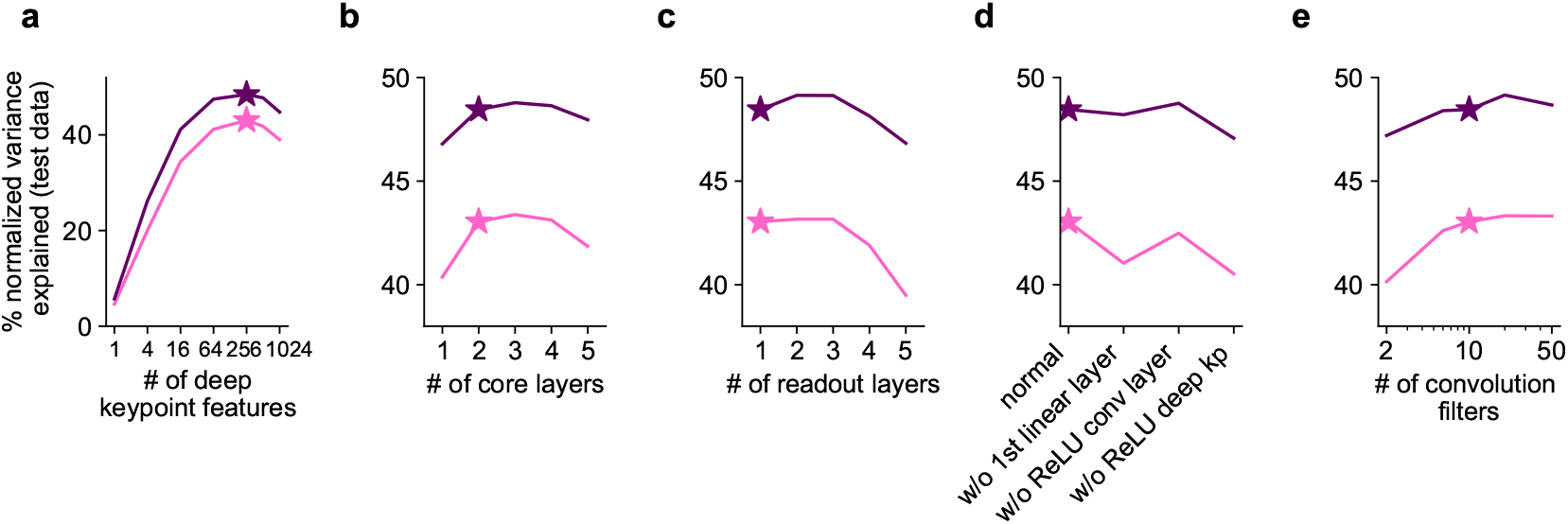
Varying the keypoints to neural activity prediction model architecture. We varied different components of the deep keypoint model and computed the normalized variance explained across neurons, choosing the architecture denoted with the star. Pink represents the average across visual recordings (n=10 recordings, 7 mice), and purple represents the average across sensorimotor recordings (n=6 recordings, 5 mice). **a**, Varying the number of units in the deep keypoint features layer – the last fully-connected layer before the output layer. Star denotes 256 units. **b**, Varying the number of core layers – the layers before and including the deep keypoint features layer, star denotes 2 layers. **c**, Varying the number of readout layers – the layers after the deep keypoint features layer, star denotes 1 layer. **d**, The performance when removing the first linear layer in the network, removing the ReLU non-linearity in the convolution layer, or removing the ReLU non-linearity in the deep keypoint feature layer. **e**, Varying the number of one-dimensional convolution filters, star denotes 10.

**S4:**
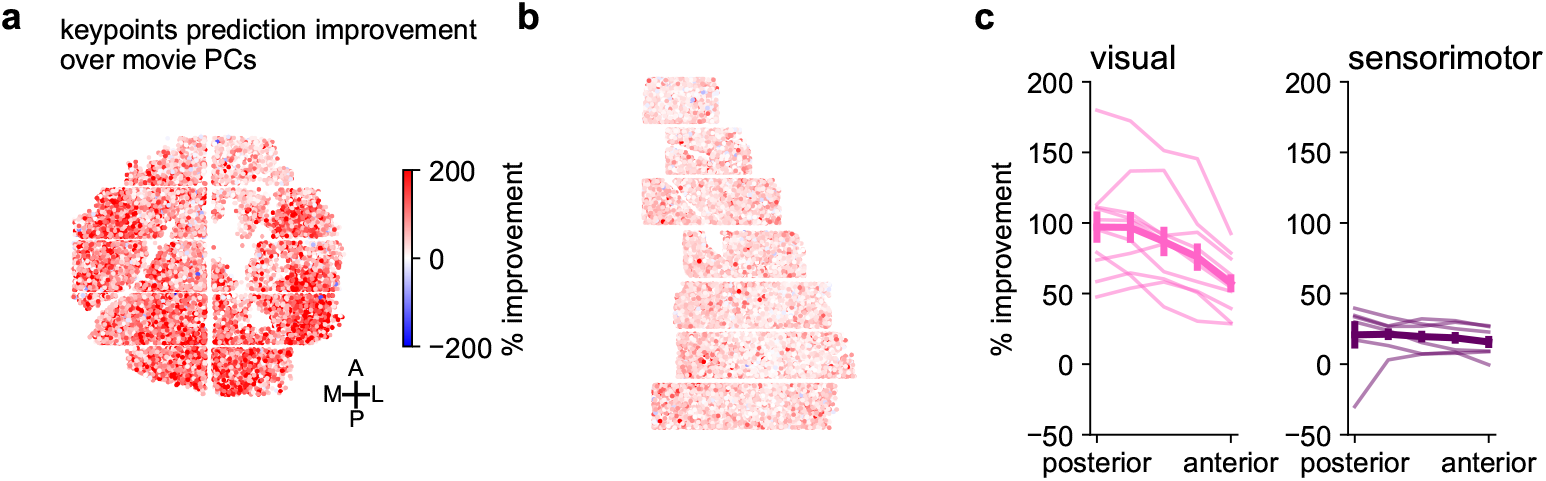
Improvement of keypoints prediction performance over movie PC prediction performance largest in posterior brain regions. **a**, Percentage increase in performance of neural prediction from keypoints over movie PCs per neuron in the example visual recording shown in Figure 3d. **b**, Same as **a** for the example sensorimotor recording shown in Figure 3e. **c**, In each recording, neurons were placed in 5 quintiles according to their position, with the leftmost point being the most posterior quintile, and the rightmost point being the most anterior quintile. The percentage improvement was averaged across neurons in each quintile, shown for each recording (thin lines), and averaged across recordings (thick line). Error bars represent s.e.m., n=10 recordings in 7 mice. **d**, Same as **c**, for sensorimotor recordings. Error bars represent s.e.m., n=6 recordings in 5 mice.

**S5:**
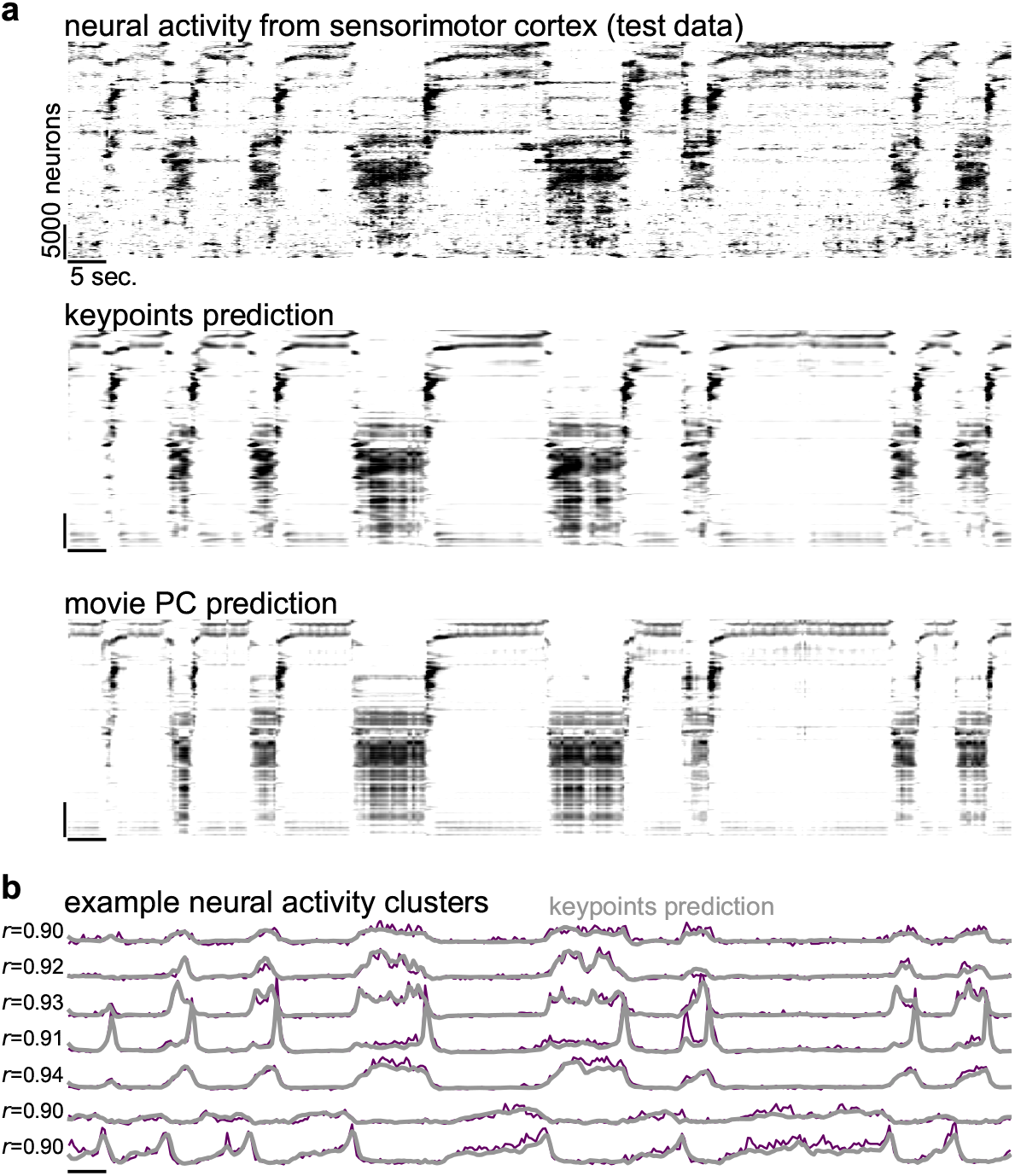
Activity in an example sensorimotor recording. **a**, Activity of the sensorimotor recording during a test period shown in Figure 3e and Figure 4c, sorted by Rastermap, along with neural predictions from the deep keypoint model and the movie PCs. **b**, Example neural activity clusters from the recording (purple), plotted with the prediction from keypoints in gray.

**S6:**
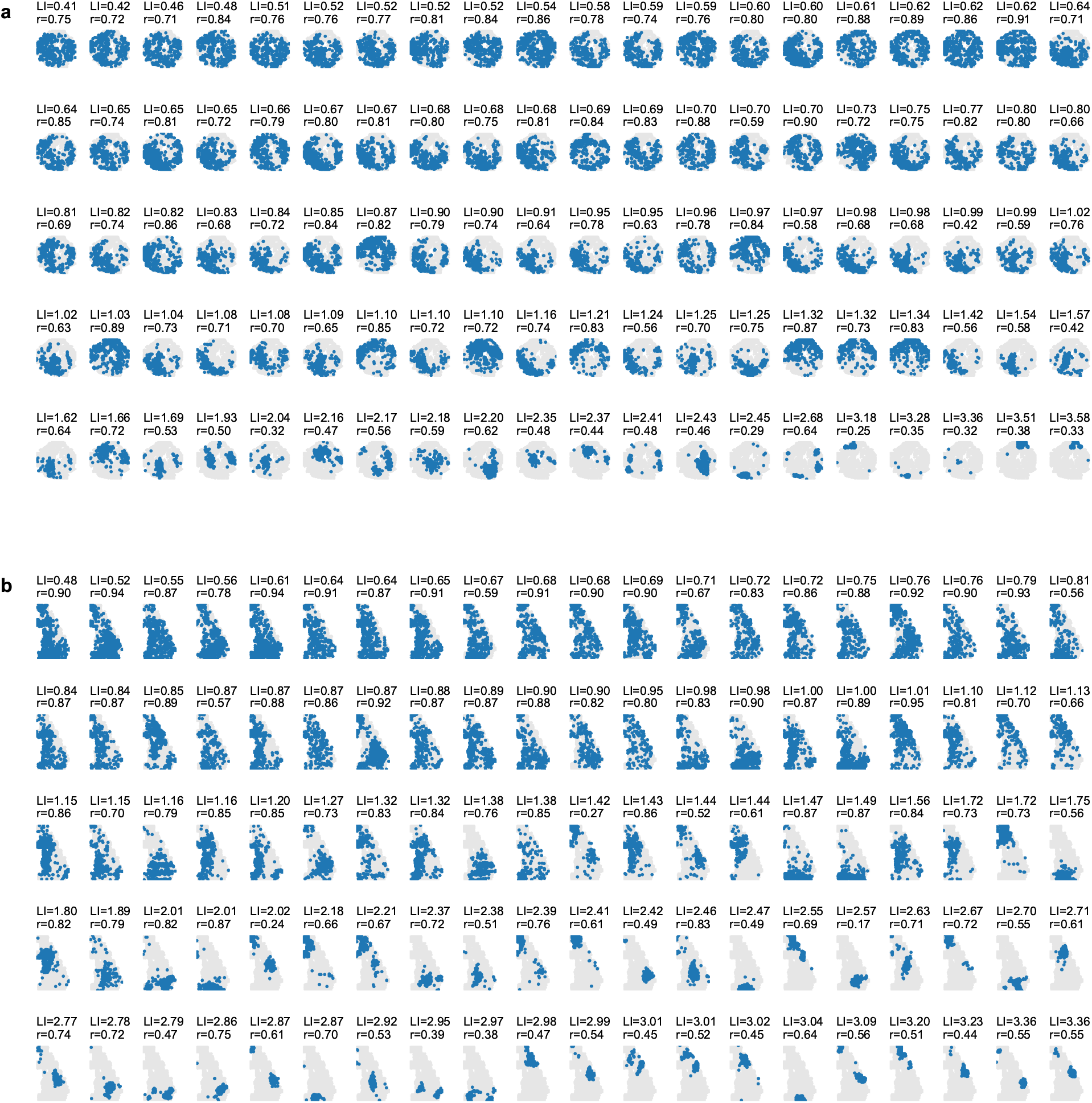
Neural activity clusters from a visual and a sensorimotor recording. **a**, The spatial locations of neurons from each neural activity cluster from the recording shown in Figure 3d and Figure 4a,b. Blue indicates neurons in the cluster, gray indicates all other neurons. LI = locality index, *r* = correlation with behavior prediction on test data **b**, Same as **a** for the neural activity clusters from the sensorimotor recording shown in Figure 3d, Figure 4c, and Figure S5.

**Supplementary video 1: Mouse face video showing keypoint tracking of the test mouse**.

**Supplementary videos 2: Mouse face videos and keypoint tracking from other lab data (Figure 2) using the base model, and the scratch and fine-tuned models trained with 10 frames**.

**Supplementary Videos 3: Visualization of HMM states**. Visualization of 4 example states from the HMM shown in Figure 5, with 15 instances of each state shown. Up to 5 seconds of time is shown in total at a speed of 0.5x realtime, with 0.5 seconds before the state starts shown (fade in from gray), and 1.0 seconds after the state ends is shown, if the state is less than 3.5 seconds (fade out from gray).

Videos available at https://drive.google.com/drive/folders/1j3Fo1nlxMZGI-XeoHSG1Wq_QSigsSog2?usp=share_link

